# Human Group Size Puzzle: Why It Is Odd That We Live in Large Societies

**DOI:** 10.1101/2022.02.18.481060

**Authors:** Tamas David-Barrett

**Affiliations:** Trinity College, Broad Street, Oxford, OX1 3BH, UK

**Keywords:** coordination, behavioural synchrony, group size, agent-based model, social networks, social technologies

## Abstract

Human groups tend to be much larger than those of non-human primates. This is a puzzle. When ecological factors do not limit primate group size, the problem of coordination creates an upper threshold even when cooperation is guaranteed. This paper offers a simple model of group coordination towards behavioural synchrony to spell out the mechanics of group size limits, and thus show why it is odd that humans live in large societies. The findings suggest that many of our species’ evolved social behaviours and culturally-maintained social technologies emerged as a solution to this problem.

## Introduction

Humans, like all non-human apes, and most non-ape primates, evolved to form social groups [1, 2]. Despite the similarity in many of traits of social behaviour, human groups differ from non-human primate groups in several features, one which is their size [3, 4]. Yet, this observation comes with difficulties.

There have been many attempts to assess the ‘natural’ or ‘ancestral’ group size of humans, often based on forager cultures today, or the – admittedly dotted – archaeological record [5–7]. One difficulty is that ecological setting rather than any inherent feature of social dynamics is often the main driver of forager group size [8–10].

The second difficulty is methodological, the core of which is the definition of a ‘human group’. Hunter-gatherers all across the world self-organise into hierarchical group macro-structure [11–13], which is more in line with a baboon-like troop organisation than a bonobo or chimpanzee one. This is despite the fact that our micro-organisation is very different from baboons, the latter being a clan dominated by a single male, while we form multi-male-multi-female societies in which the smallest unit is predominantly pair-bonded, the human equivalents of the baboon ‘clan’ [14].

Yet, even given these difficulties, it is clear that the natural human group [15, 16], at the average of 840 individuals (defined as the highest level of the social organisation, measured in 340 forager cultures [17]), is much larger than the largest coherent baboon bands at 220 individuals [18] or chimpanzee groups that are typically around 40-45 individuals (but, can reach up to 150-200, even if rarely) and is the largest of any ape group [19, 20]. (NB. Baboon bands sometimes form large meta-populations, but these are more similar to herds than groups in the ‘ape group’ sense.) Thus, independent whether ecology drives the presence of relatively small human group, the maximum size of the ‘natural’ human group is much larger than of other apes’, and of non-ape primates.

Furthermore, in different technological environments, humans regularly form groups of millions and even billions of people. The median country size today is 6.7mn people, and the median size of the largest cities in our countries is 2.1mn people [21]. Arguably, the global society is so interconnected [22, 23] that it can be regarded as a group itself, despite its improbably large size, of currently 7.8bn people. (It is not clear, however, it is justified to count modern societies as natural groups. We know from other areas of human life that modernity can change our ancestral ‘setting’, for instance the majority of us gave up foraging for farming [24, 25], equity for inequity [26, 27], and kinship-based social networks for friendship-based ones [28, 29].) Thus, not only the ‘natural’ group size is so much larger than the groups of non-human primates, but our modern societies are so enormous that there may not be a natural group size to start with.

This is a puzzle. Humans are apes, and as such also primates. How come that our species’ groups are so big? And if our groups *can* be this large, how come that other apes’ and non-ape primates’ groups are not as large? Much of the scientific literature about human social evolution has been concerned with the tricks and ruses our species evolved and invented to allow a large group size. But under these shelves of papers lies the assumption that forming super large groups is something unnatural, something odd. If there is no limit, why would there be a need for a trick to break through it? To my knowledge, there is no paper in the literature that spells out this assumption by asking the question: why is it odd that we live in large societies?

This paper explores this question through a simple model of the network dynamics, and aims to illustrate one particular group size constraint type: coordination limits.

## Methods and Results

Let us take *n* agents that form a randomly connected graph of degree *k*. These agents face a problem, for instance, set by the ecological niche they live in, such that they must coordinate their action to be able to act as one.

The coordination takes place on a unit circle, as if each task was finding a shared direction on a compass [30–33].

The agents start with a randomly assigned initial value draw from a uniform distribution on a compass:

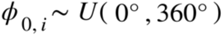

For the group to reach behavioural synchrony, the agents go through a series of pair-wise meetings in which they synchronize their *ϕ* values. For a meeting, or synchronisation event, two agents are randomly picked, and set their *ϕ* values to the mid-point of their old *ϕ* values:

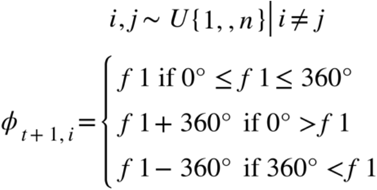

where

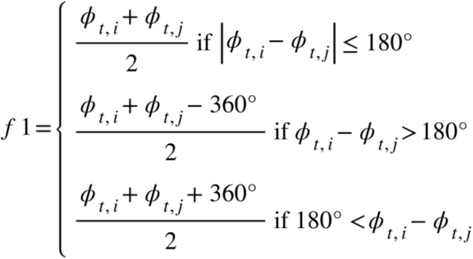

Sidenote: using compass coordination might seem a little cumbersome merely to go to the middle of some range, however, the choice of this subject of synchronisation is not arbitrary. Unlike in this case, when the initial values are on a finite linear range, the outcome of the synchronisation has Gaussian normal distribution even if the initial values are uniformly distributed. However, when the values are on a unit circle, i.e., they represent compass directions, the outcome of the synchronisation is uniformly distributed if the initial values are also uniformly distributed (Fig. 1). As a consequence, in the latter case the final point cannot be guessed, which is useful for models in which the agents become smart with memory and foresight.

**Fig. 1.**
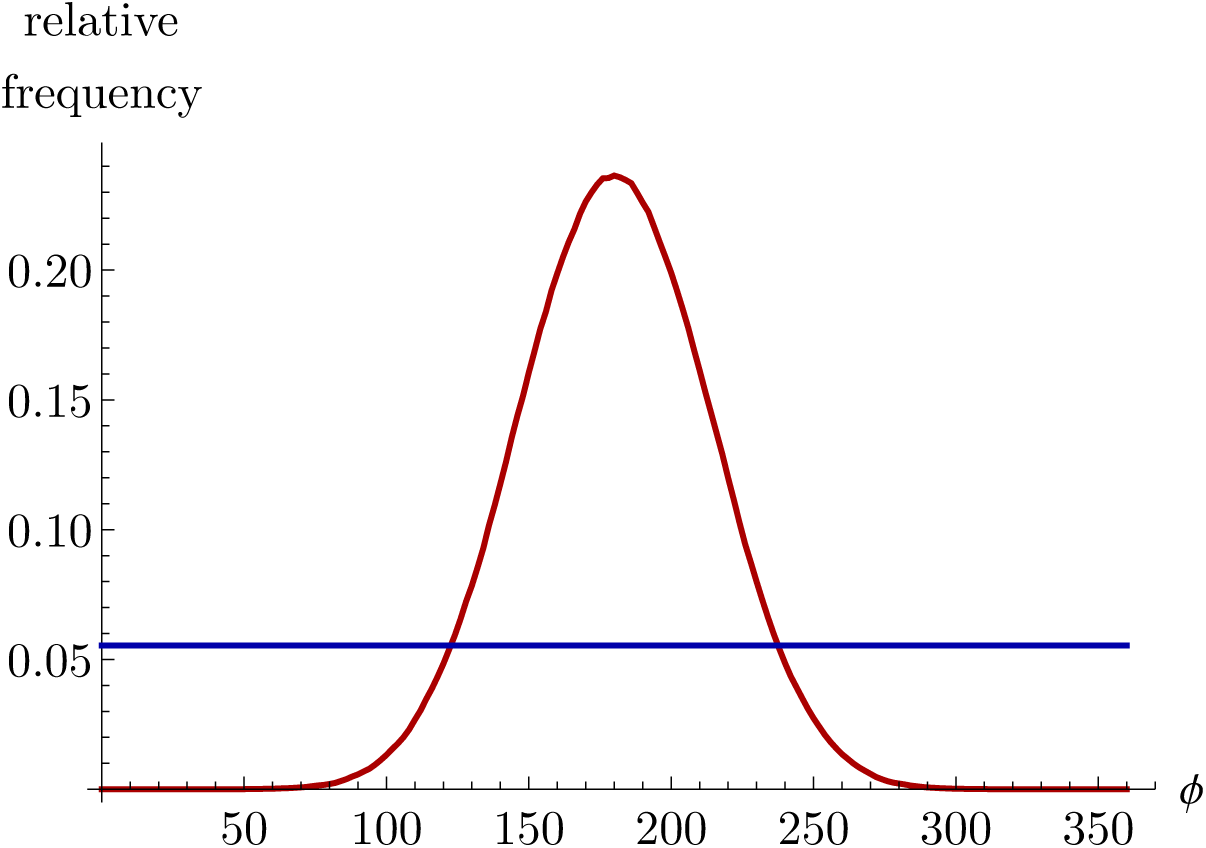
Network coordination outcome distributions when the synchronisation is on a linear range (red line) vs. a compass direction (blue line), both cases starting from uniform distribution (which is identical to the blue line).

Let *δ* denote the average distance among the *ϕ*s of all the group members.

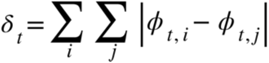

where *t* is the average number of meetings an agent has, and the | | distance measure refers to the nearer side of the unit circle. (For instance, the distance between 10 and 90 is 80, while the distance between 10 and 350 is 20.) Notice that *δ* is, an inverse measure of synchrony, i.e., group’s coordination efficiency is high when *δ* is low, and vice versa.

Independent of network structure, as long as k>2 and the graph is connected, the *ϕ* values converge and the *δ* goes to zero with *t* increasing (Fig. 2). That is, as the number of meetings goes up, the group turns towards the same direction.

**Fig. 2.**
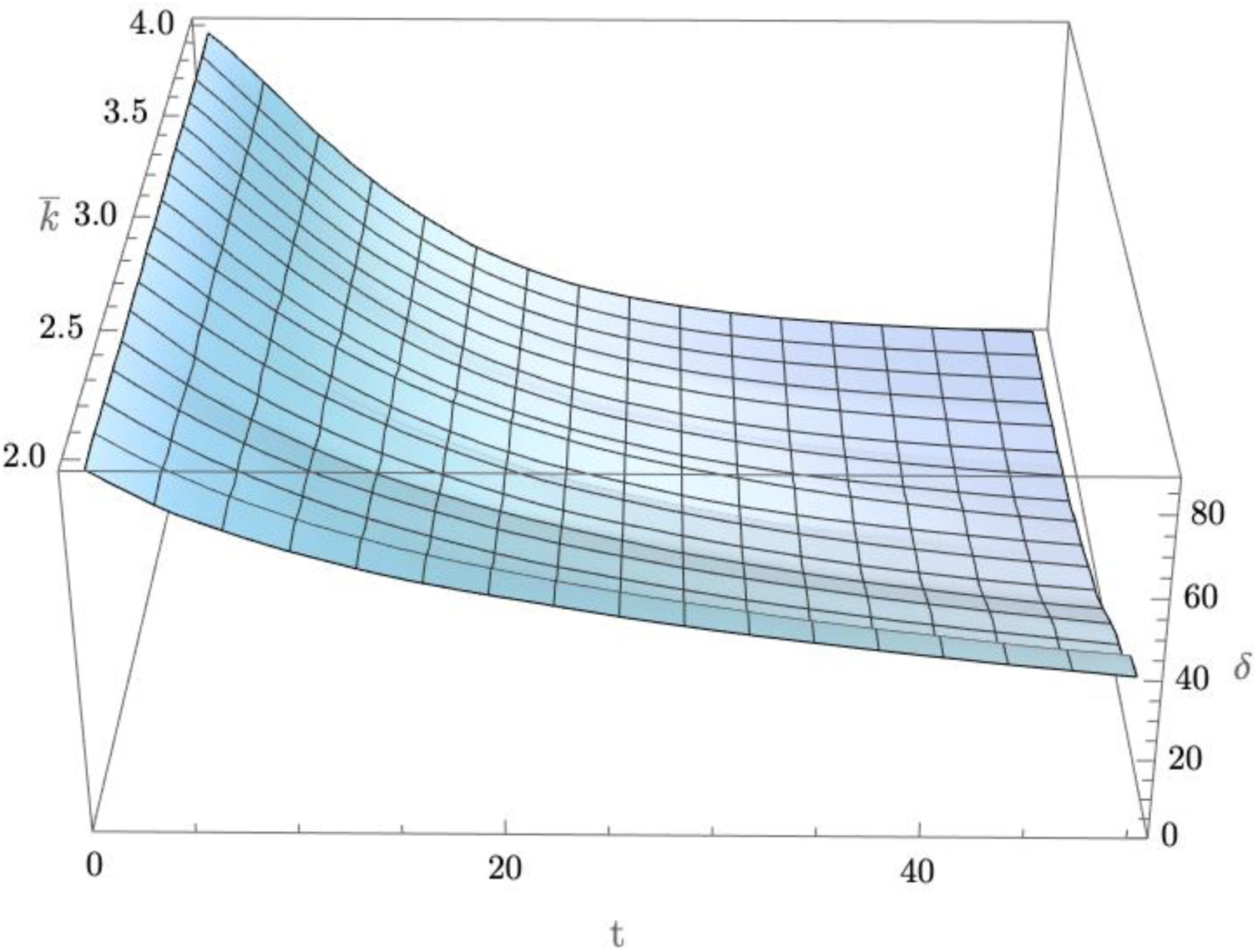
The compass direction converges among the group members as they go through a series of dyadic synchronisation (the horizontal lines all go down in delta), and the convergence is faster, the higher the average degree is (the vertical lines also go down in delta). The x-axis: time, i.e., the average number of meetings among the group members, y-axis: the average degree of the graph, i.e., number of connections per agent, z-axis: the average distance in compass direction among the agents.

### This pattern of convergence is not surprising

This is an instantiation of an established result in network science, often employed to describe innovation diffusion in social networks [34, 35]: unless k=2 *and* the graph is circular, convergence happens. The only question is the speed.

Notice that the speed of convergence is driven by the average number of edges per nodes (Fig. 2). This is also not surprising: the more connected a network is, the faster the synchronisation is, another established network science fact. This speed difference is important, because if the network describes a real-world human (or other species’ cooperating) group of individuals, then the amount of time that the group members can spend is likely to be limited by ecological, technological factors.

Let us introduce a time limit, denoted by *τ*, as a parameter external to the coordination problem (Fig. 3). Not surprisingly, given any arbitrary time limit, the higher the degree is, the more synchronised a group is at the cut-off point.

**Fig. 3.**
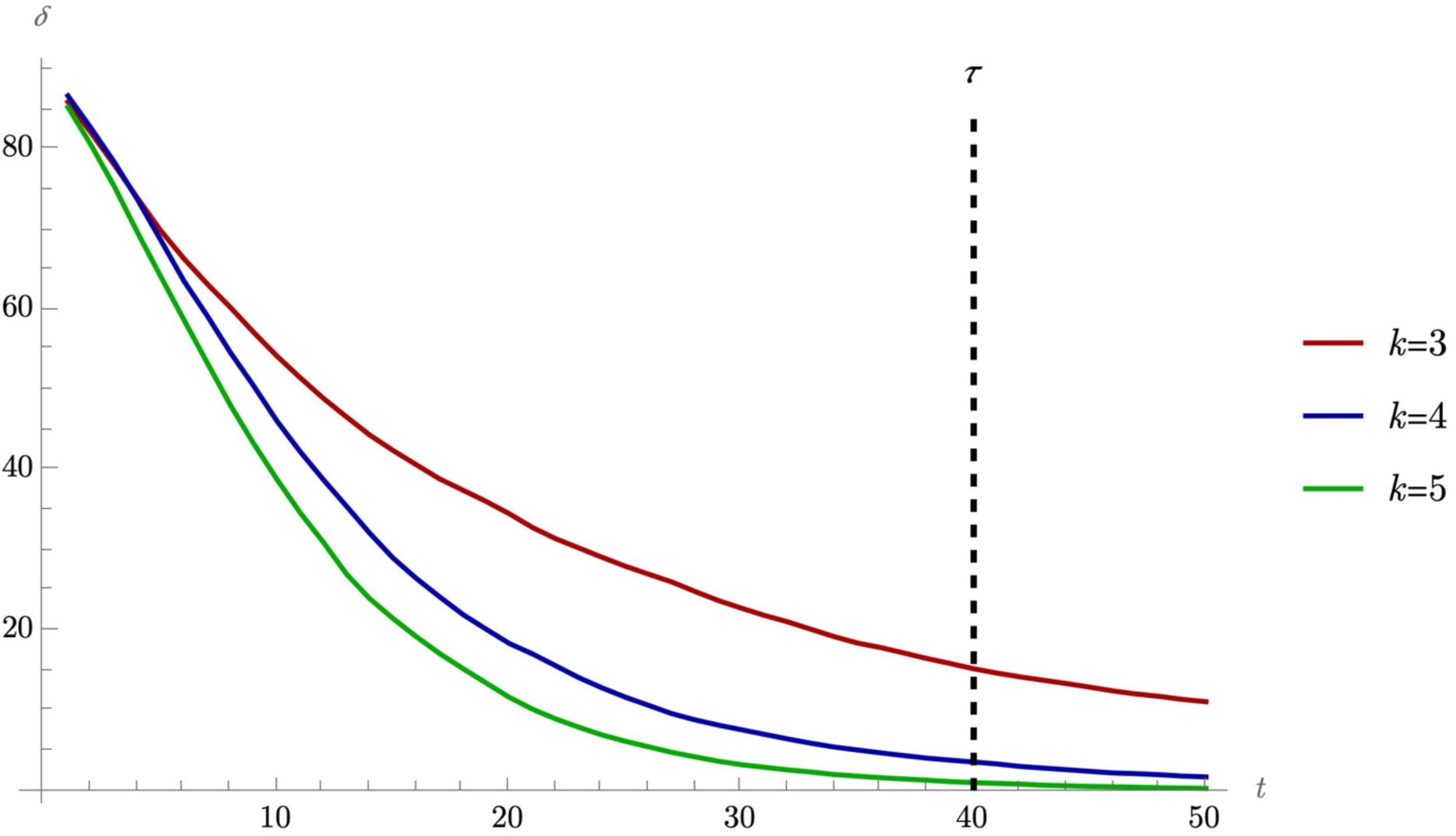
Introducing a limit to the number of meetings the agents can have illustrates the importance of the fact that the speed of convergence varies with the average degree. (*n*=20.)

Thus, the time limit to coordinating the action, a constraint that all human and other animal groups face in practice, makes the degree of the social network play a key role. This raises the question of degrees origin. Where does the number of connections per agent comes from?

Let us assume that evolution (or economics) works on the level of the individuals, and thus that the agents decide their number social connections for themselves. Let us assume that social connections come with benefits and costs that translate into a payoff:

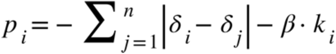

where *p* is payoff or evolutionary fitness, and *β* is the cost parameter.

To evolve the optimal degree, let us use an evolutionary algorithm the following way:

Step 1: For a group of *n* agents, assign random degree uniformly to each agent such that *k*_*i*_ denote the degree of node *i*, with one agent having *k*-1, and one *k*+1 connections. (This setup ensures that the corresponding graph exists.)

Step 2: Run a large number of group coordination events, with a fixed limit of average number of meetings at *τ*. Calculate the mean payoff for each agent.

Step 3: Given the payoffs, set *k* to the degree number of the best performing agent. Repeat steps 1-3 for an evolutionary round.

Using this selection algorithm, the degree evolves in line with the social costs of an edge (Fig. 4). If contact maintenance costs are low, then the individuals are best off to have many social connections, and vice versa, when costs are high, the social network will be sparse.

**Fig. 4.**
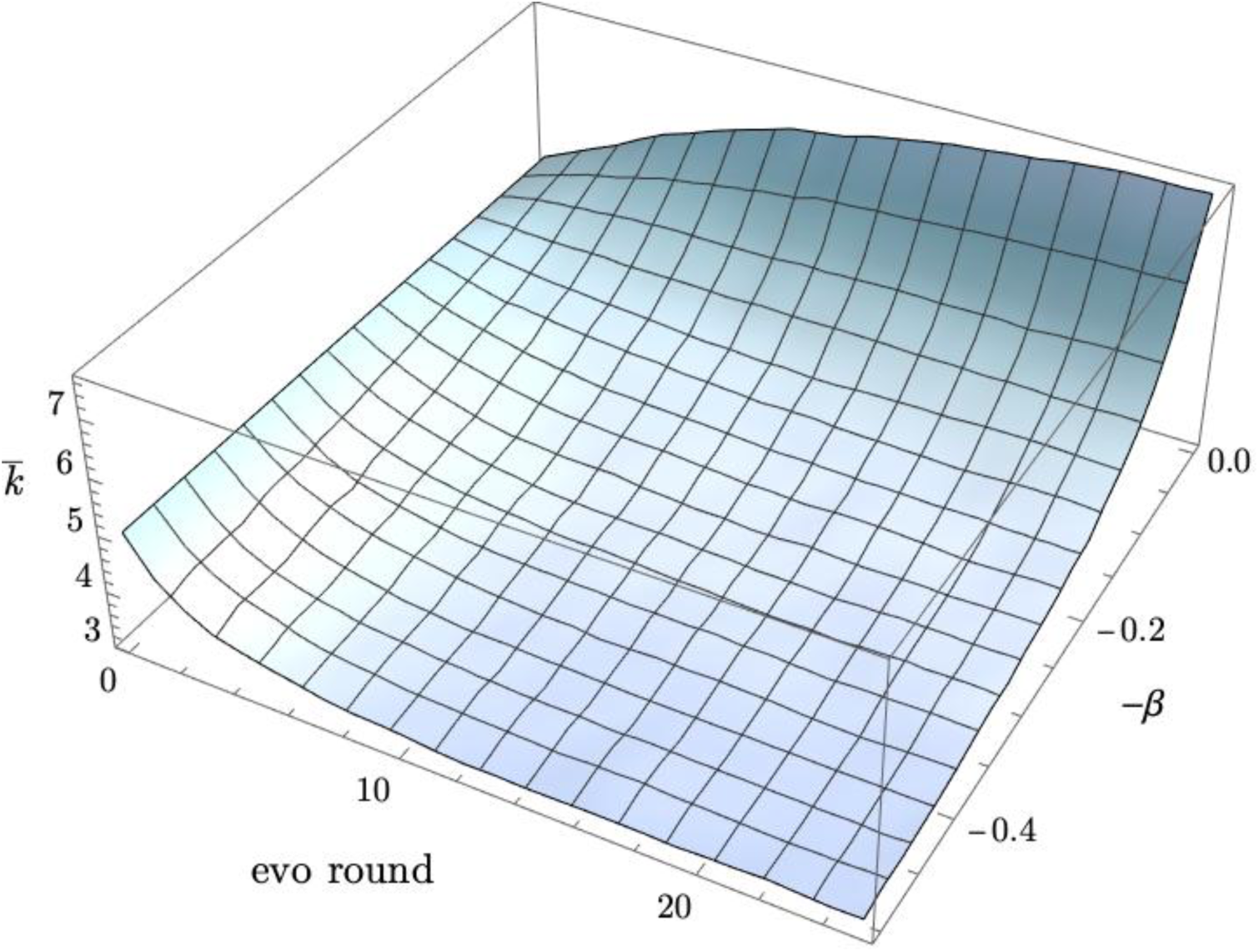
Evolving the number of social contacts as a function of the cost of maintaining social relationships. The x-axis: evolutionary round, y-axis: the cost of maintaining a single relationship, and z-axis: the average degree among the agents. (*n*=20.)

Thus, from the fact that the group needs to coordinate its action comes two, partially-opposing interests. On one hand, the group’s coordination efficiency increases when there are more connections among the agents (Fig. 2). On the other hand, the agents will end up limiting the number of connections if these are costly (Fig. 4).

Notice that there are two kinds of costs in this model one on the individual’s level, and one on the group’s level. The cost of an edge is suffered by the individual, and is not linked to the group’s coordination problem. However, there is a link between the two costs as a loss of group-level coordination efficiency emerges if of *k* less than maximum.

Notice also that the given the coordination problem, the two parameters *τ* and *β* together determine the relationship between group size and synchronisation efficiency (Fig. 5).

**Fig. 5.**
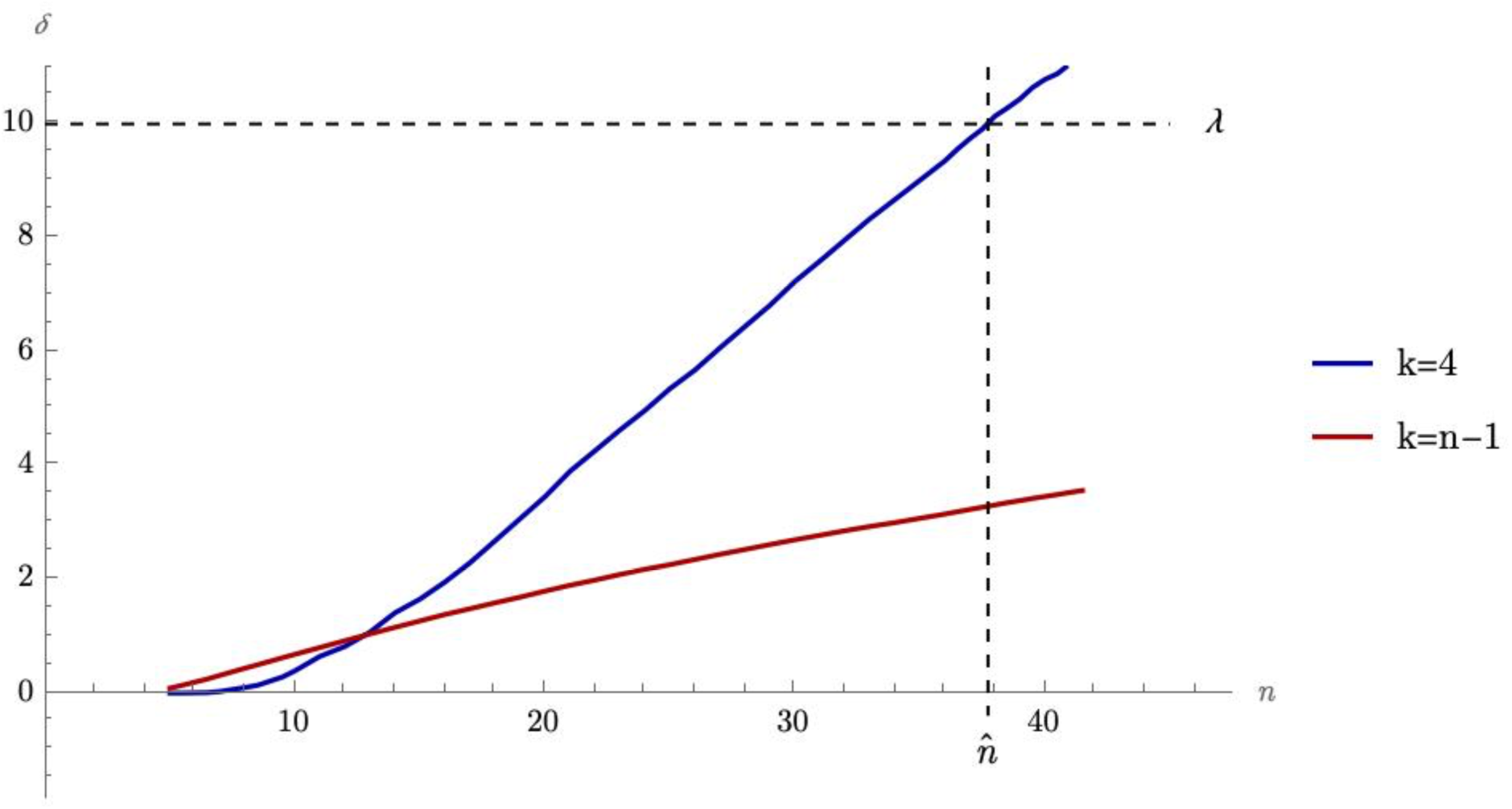
A minimum coordination efficiency threshold also limits the maximum group size. (*n*=20.)

If the ecological or technological problem the group faces is such that there is a coordination threshold above which the group action falls apart, and under which the group operates as a unit, then this limit also serves as a maximum group size limit. Let *λ* denote such a threshold in *δ* then given the relationship between n and *δ*, the maximum group size is set, 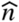.

Notice that as the *δ*(*n*) function is entirely determined by *τ* and *β*, and we get 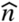 from 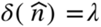, it is the three parameters and nothing else that determines the maximum group size. Thus, it is the ecological setting together with the nature of group coordination that set the maximum group size. Above this, a group cannot function in this model.

## Discussion

This paper used three parameters as a frame to the collective action coordination: time limit, cost of having a friend, and minimum coordination efficiency threshold. The results showed that in a behavioural synchrony framing these three parameters together determine a maximum group size.

There are well documented human behaviours in which the need to reach behavioural synchrony efficiently limits group size. For instance, the result sheds light on why jazz jam sessions have only small band size [36–38].

Thus, three empirical questions follow: (a) is there should a group size limit for non-human primates, and (b) if so, do human groups break out of this limit, and (c) and if so, are there special evolved or cultural ‘solutions’ in place to facilitate going through such a threshold

First, the group size distribution of non-human primates (Fig. 6) is in line with the models suggestion that such maximum group size exists. Our closest relatives’, the other great apes’, average group size range ranges from 9 (eastern gorilla), to 42-46 (chimpanzee and bonobo), with all the other orangutan and gorilla species falling towards the lower end of the range [19, 39, 40].

**Fig. 6.**
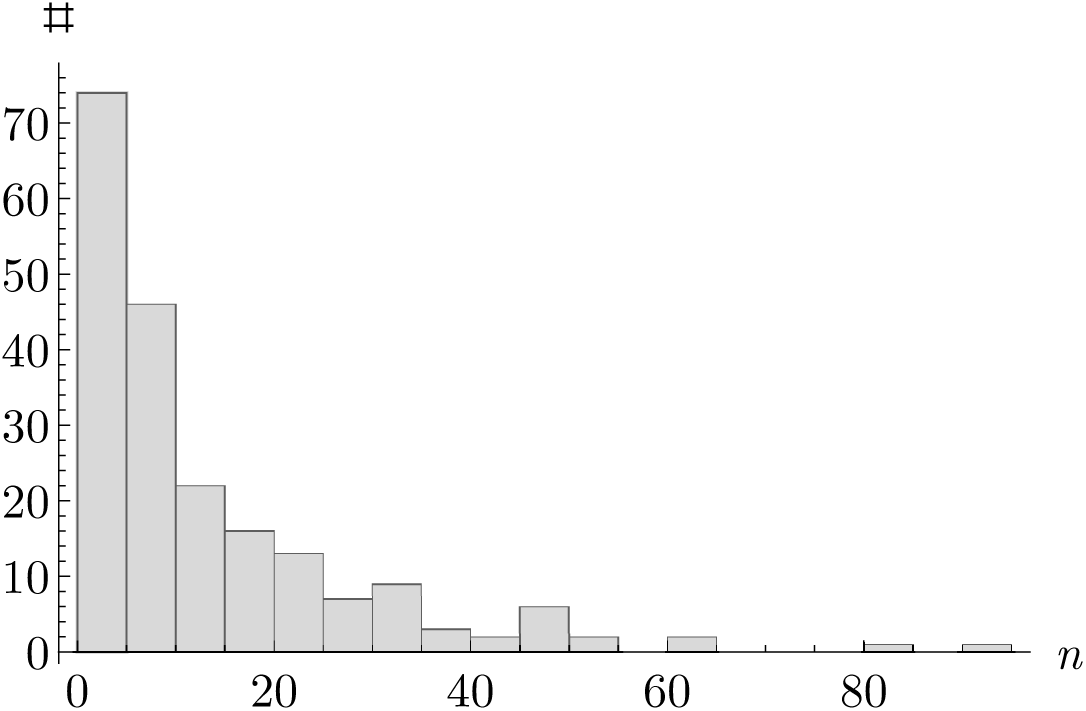
Non-human primate group size distribution, where x-axis: group size, y-axis: number of primate species with *n* being their average group size. [19, 39–50])

Second, ethnographic data is in line with the suggestion that the human group size is beyond a threshold that other primates did not cross.

For instance, let us consider the case of small languages, which is most of them: the median speaker number of a language is under 1000 on all continents [51, 52]. The highest language diversity on the planet is on the island of New Guinea, which consists of Irian Jaya, which belongs to Indonesia, and the mainland of country of Papua New Guinea. The extremely high language diversity is due to the geography of the island. As the Australian continental plate pushed northward against an easter protrusion of the Pacific plate [53], the crust of the Earth formed a mountain range with hundreds of deep valleys, almost every one corresponding to a different language [54].

In this highland region, crossing other group’s territory is highly perilous, intergroup violence is ongoing. (For instance, in 2008, the director of the of the Timika hospital told me that one of the most frequent injury they had to deal with was from arrow wounds. Similarly, when I asked tribal leaders about with whom they had conflict, they invariably mentioned nearby, competing tribes rather than the Indonesian military.) Thus, the most likely explanation for the separation of the languages has to do with the fact that it is impossible to maintain military domination across several valleys at the same time. As an exception that supports the point, the Dani culture, which has approximately 600,000 speakers, occupies the only large flat area of Irian Jaya Highlands.

Thus, the median number of speakers per language, which is 643, likely corresponds not-only to an ethnolinguistic meta-population, but also to a reasonable estimate of the number of people who live in a relatively small valley, relying on each other for protection against the enemies around. For this reason, 643 may be seen as a good estimate for what the highest-level group size in the Highlands, whether they split into separate settlements or not. This is in line with the observation from a very early, 1961, population survey that while the average settlement size was 159-170 people, depending on measurement [55, 56], the settlement sizes were driven by ecological factors, with the largest reaching above 1000 people [54]. (NB. The Irian Jaya, i.e., the Western half of the island of New Guinea, was first entered by outsiders in the mid 1950s, and hence the Highlands part of the 1961 survey must have covered stone-tool technology using cultures that lacked the concept of metal or pottery.)

Thus, using the New Guinea example, the average human average group size is at least 4 times as high as that of the great apes, with possibly 15 times larger. And when comparing maximum group size, the difference is even starker. The largest non-human great ape group ever recorded was the Ngogo chimpanzee ‘community’ that ranged between 140 and 206 members [57]. Compare that to the 1000+ settlement size of the 1961’s New Guinea maximum settlement size, let alone human communities today that go into the tens of thousands to millions.

Third, how is this large human group size possible? To what extent did our species tricks in ‘social technologies’ evolve or were invented to solve the problem of the looming group size limit? Did language evolve as a third-party information tool to facilitate larger groups [58–60]? Did the structural solutions for macro-network management, like fission-fusion pattern, evolve to facilitate large groups or to facilitate temporal variation in the environment’s carrying capacity [61–65]? In particular, is baboon-like fission-fusion dynamic a clue towards the evolution of the ability to form large complex groups in humans, or is this a case for parallel evolution [14, 66]? Is a central figure’s one-way communication, as in the case of priesthood [67], or in social technologies facilitated by mass media [68, 69], another social technology that allows larger groups?

(Note that this paper focuses on group size being limited only by coordination efficiency. This model assumes away the problem of cooperative stance, and dyadic cooperation is implied in the cost of maintaining the social network edges. Yet, there is an entire library on the origins of costly cooperation [70–77], and the interaction between network structure and cooperative stance [78–99], and even the possible conflict between the dynamics [100–102], represented by the negative relationship between the social cost parameter of this paper, and the maximum group size. To the extent the interaction between two coordination and cooperation dynamics shaped the evolution of network building traits is subject of future research.)

## Notes

### Competing Interest Statement

The authors have declared no competing interest.

